# Arbuscular mycorrhizal fungal diversity and association networks in African tropical rainforest trees

**DOI:** 10.1101/2024.02.06.578868

**Authors:** Damilola Olanipon, Margaux Boeraeve, Hans Jacquemyn

## Abstract

Tropical rainforests represent one of the most diverse and productive ecosystems on Earth. High productivity is sustained by efficient and rapid cycling of nutrients through decomposing organic matter, which is for a large part made possible by symbiotic associations between plants and mycorrhizal fungi. In this association, an individual plant typically associates simultaneously with multiple fungi and the fungus associates with multiple plants, creating complex networks between fungi and plants. However, there are still very few studies that have investigated mycorrhizal fungal composition and diversity in tropical rainforest trees, particularly in Africa, and assessed the structure of the network of associations between fungi and rainforest trees. In this study, we collected root and rhizosphere soil samples from Ise Forest Reserve (Southwest Nigeria), and employed a metabarcoding approach to identify the dominant arbuscular mycorrhizal (AM) fungal taxa associating with ten co-occurring tree species and to assess variation in AM communities. Network analysis was used to elucidate the architecture of the network of associations between fungi and tree species. A total of 194 AM fungal Operational Taxonomic Units (OTUs) belonging to six families were identified, with 68% of all OTUs belonging to Glomeraceae. While AM fungal diversity did not differ between tree species, AM fungal community composition did. Network analyses showed that the network of associations was not significantly nested and showed a relatively low level of specialization (*H*_2_ = 0.43) and modularity (*M* = 0.44). We conclude that, although there were some differences in AM fungal community composition, the studied tree species associate with a large number of AM fungi. Similarly, most AM fungi had a large host breadth and connected most tree species to each other, thereby potentially working as interaction network hubs.

**Graphical Abstract:** 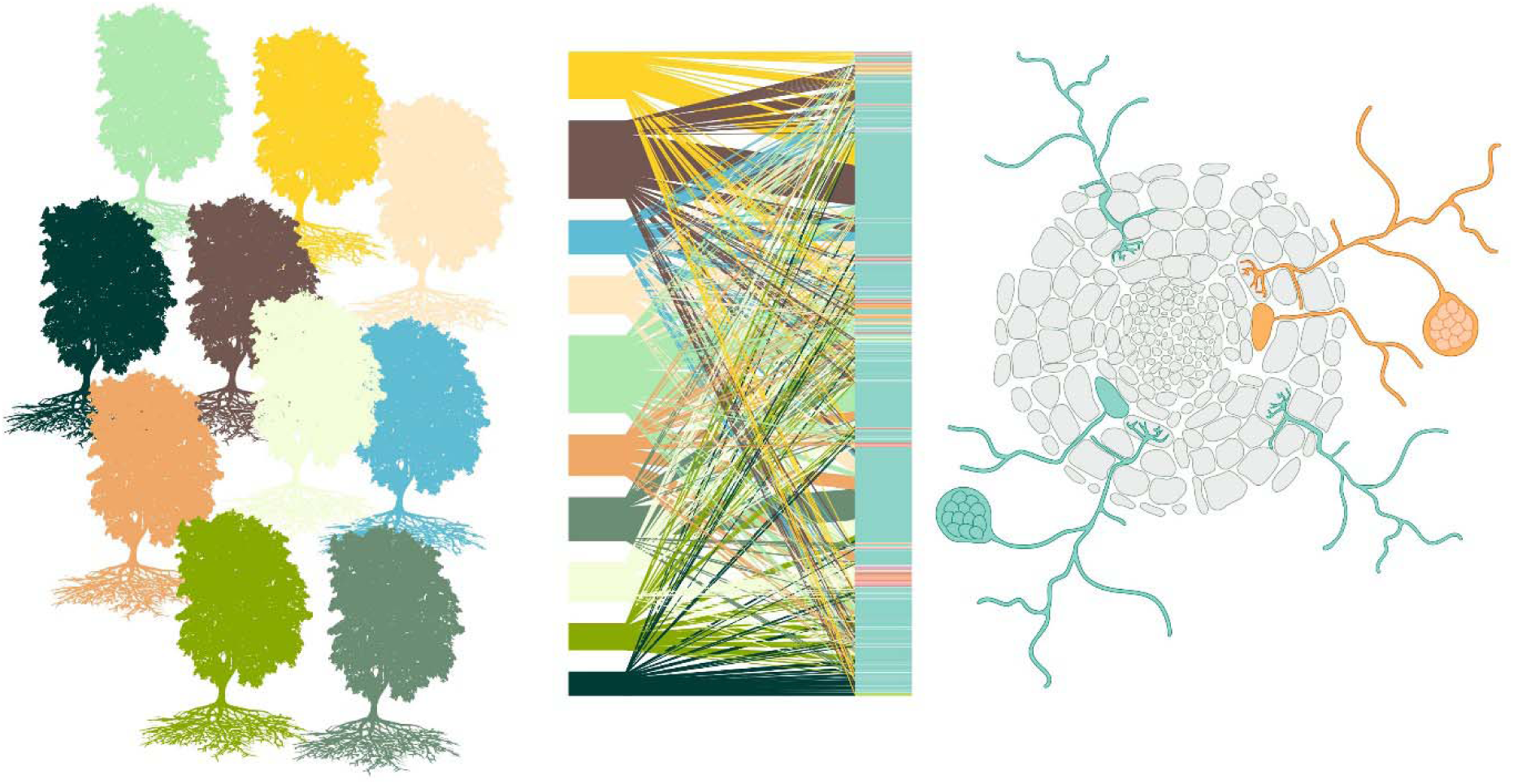

## Introduction

Tropical rainforests represent one of the most diverse ecosystems on Earth, containing over half of the Earth’s 5 to 20 million species (Gibson et al. 2011; Mora et al. 2012; Scheffers et al. 2012). Trees are the dominant growth form of tropical rainforests and account for most of the plant diversity (up to 42%) (Taylor et al. 2023). Tropical tree communities are typically characterized by high species richness, often surpassing 100 species per hectare (Losos and Leigh 2004), and in some cases >300 tree species per hectare have been recorded (Gentry 1988). Besides, tropical rainforests are characterized by an enormous productivity and biomass (Whittaker and Likens 1975), notwithstanding the highly infertile soils on which they often occur. Apart from year-round water availability and warm temperatures, this high productivity is sustained by efficient cycling of nutrients through decomposing organic matter (Vitousek 1984), which is for a large part made possible by symbiotic associations between plants and mycorrhizal fungi (Smith and Read 2008; van der Heijden et al. 2015). In this association, the fungi provide nutrients, particularly phosphorus and other mineral elements, and water to the plant and in return the plants provide carbon derived from photosynthesis to the fungi (Smith and Read 2008).

Basically, plants form four types of mycorrhizal associations – arbuscular mycorrhizas (AM), ectomycorrhizas (EcM), ericoid mycorrhizas (ErM), and orchid mycorrhizas (OrM) – but only two (AM and EcM) associates with trees (Brundrett and Tedersoo 2018). The EcM lifestyle is found across the fungal phyla Ascomycota and Basidiomycota, but is restricted to particular clades of plants and have only been observed in ca. 3% of land plant species (Brundrett and Tedersoo 2018). This is in sharp contrast with AM, which have been found in over 70% of all land plants. Most AM are formed by fungi belonging to the subphylum Glomeromycotina (van der Heijden et al. 2015; Spatafora et al. 2017). Although the majority of tropical trees depend on AM fungi for growth and survival (Onguene and Kuyper 2001; Alexander and Lee, 2005), most research into AM fungi has so far predominantly been done in Europe, North America and China (Větrovský et al. 2023), while mycorrhizas in tropical areas have received less attention (Alexander and Selosse 2009).

Inside the tree roots, AM fungi form arbuscules, where nutrient exchange takes place, and hyphae, which extend into the soil and are responsible for nutrient uptake and transport (Perotto and Balestrini 2023). An individual plant typically associates simultaneously with multiple AM fungi and an AM fungus often associates simultaneously with multiple plants (Van der Heijden et al. 2015; Davison et al. 2016). This creates underground fungal networks that do not only link individuals of the same species, but also of different species (Montesinos-Navaro et al. 2012; Chagnon et al. 2012). Although accurate description of these networks does not necessarily provide insights into whether these trees are truly interconnected and exchange nutrients (Karst et al. 2023), it does provide important information on how trees share mycorrhizal fungi and therefore are relevant for understanding ecological and evolutionary dynamics in species rich communities. Due to their apparent low diversity in comparison to their widespread occurrence across land plants, AM fungi have long been considered to show little specificity (Smith and Read, 2008). However, with the increased use of molecular methods, studies have found indications of specificity or preference in AM fungus-plant interactions (Husband et al. 2002; Montesinos-Navaro et al. 2012). Network analyses are highly useful to study potential interactions and preferential partner selection between plants and fungi and how these are affected by global change (Toju et al. 2015; Bennett et al. 2019; Chagnon et al. 2020).

In species-rich forest ecosystems, the networks of interactions between fungi and trees can be complex and differ from those typically observed in other mutualistic systems such as the interactions between plants and pollinators (Toju et al. 2014; 2015). Most notably, most plant-fungus networks in forest ecosystems have been shown to lack a typical nested architecture, but instead showed an “antinested” topology (Toju et al. 2015). Besides, plants and fungi were associated with a narrower range of partners than expected under null models that assumed random associations between hosts and symbionts and networks were compartmentalized into modules of closely associated plants and fungi (Toju et al. 2015). However, there are still few studies that have used a DNA barcoding approach through next-generation sequencing to identify the mycorrhizal fungi associating with trees in tropical forest ecosystems, particularly in Africa, and to assess the network of associations between trees and their fungi.

In this study, we assessed AM fungal community composition and diversity of tree roots and rhizosphere samples in Ise Forest Reserve, Southwest, Nigeria, using a molecular metabarcoding protocol that employed high throughput sequencing of a fragment of the small subunit 18 SSU rRNA gene to identify AM fungal taxa to species level. Our specific research objectives were to (i) identify and characterize AM fungal diversity in the tree roots and rhizospheric soil and (ii) carry out a network analysis of AM fungal community to elucidate potential interactions and preferential partner selection. We hypothesized that (1) most tree species associate with a large diversity of AM fungi, (2) most trees show little preference and thus share a considerable number of their fungi, and as a result (3) the network of associations shows a low degree of specialization and modularity.

## Materials and Methods

### Study site

The study was conducted in Ise Forest Reserve, a tropical rainforest with a high diversity of tree species located in Southwest Nigeria (7°23.658′N, 5°24.092′E, elevation: 357 m a.s.l.) (Fig. 1). Ise Forest Reserve covers about 142 km^2^ and is located 6 km northward of Ise Ekiti Township (Orimaye et al. 2016) and bordered by a river on the west. The forest reserve is a tropical rainforest that is surrounded by farmland and human settlements. It currently faces several threats, including illegal logging activities, marijuana cultivation, and conversion to farmlands. Owing to these, the government of the state has issued an executive order establishing a conservation area within the forest reserve to protect its rich flora and fauna, and it is now termed a nature forest reserve. The study site is dominated by 50- to 100-year-old trees such as *Pycnanthus angolense, Albizia ferruginea, Celtis zenkeri, Terminalia superba, Ficus mucuso, Pseudospondias microcarpa, Triplochiton scleroxylon, Amphimas pterocarpoides, Mansonia altissma, Blighia sapida, Sterculia tragacantha, Pterygota macrocarpa, Lecaniodisus cupanoides*, and several others. Crown heights of tree species were ≥ 30 m, forming a closed tree canopy with understory populations of herbs, shrubs and lianas. The mean annual precipitation and temperature is about 1380 mm with alternating wet and dry seasons and 25.3 °C, respectively. The soil of the region can be classified as well-drained silty-loam.

**Fig. 1.**
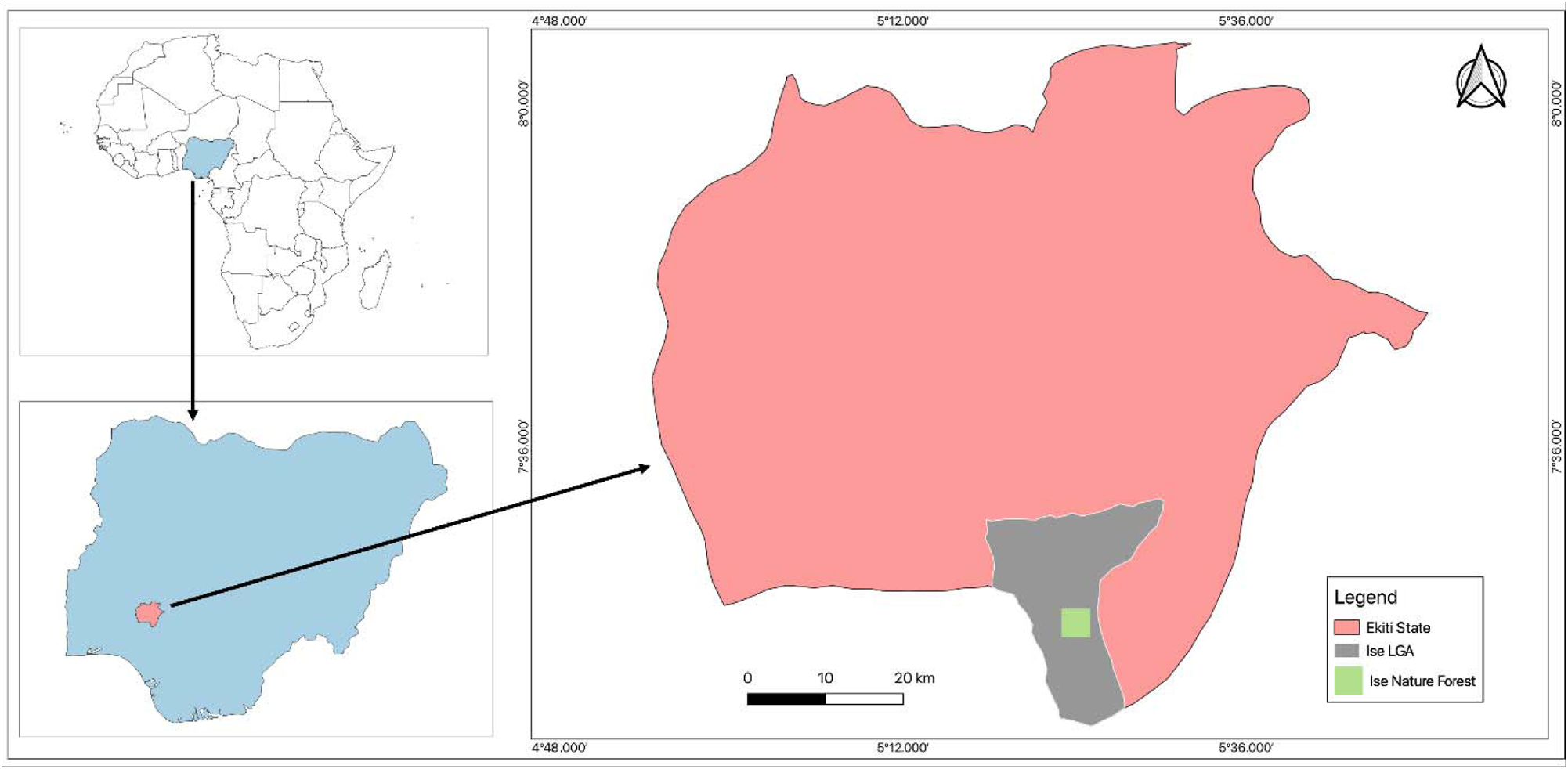
Map showing the location of the study area (Ise Forest Reserve) in Nigeria.

### Sampling

Root and soil sampling was carried out in January 2022. Standard plots of 50m x 50m were established in species-rich portions of the forest. Five stands of each tree species were sampled to have samples that are representative of the diversity within the 23 individual tree species (*n* = 115). Fine roots (1 – 3 cm; <2 mm diameter) were obtained by digging along a large root starting from the base of the tree to ensure the sampled roots were from the parent tree. Terminal roots were collected within a 2 m radius of the tree and within the upper 20 cm of the soil. Up to 20 fine root samples from the same tree were pooled in paper bags with silica gel to make a composite sample and were labelled appropriately. Core soil samples (0 – 20 cm depth) were collected from each location where a root was excavated after which they were mixed to form composite samples (*n* = 23). Soil was sieved (2 mm) to remove stones and root tissue and then divided into two fractions: one for soil chemical analyses, and the second fraction for DNA extraction. All samples (root and soil) were transported to the Laboratory of Plant Conservation and Population Ecology of the Department of Biology, KU Leuven, Belgium for further analyses.

### Soil analyses

Soil samples were analyzed for determination of pH, phosphorus, ammonium and nitrate concentration, gravimetric water content and organic matter content. Soil was mixed with deionized water in a 1:10 ratio and shaken for 10 min before measuring the pH with a pH probe. Soil phosphorus content was determined using the Olsen-P extraction and subsequent colorimetrical analysis using the molybdenum blue method (Robertson et al. 1999). Ammonium and nitrate concentrations of soil samples were measured by shaking 5 g of soil in 25 ml of 1 M Potassium Chloride (KCl) for 30 min, after which the solution was centrifuged (5 min at 3,500 g) and the obtained supernatant was analyzed colorimetrically using an Evolution 201 UV-visible spectrophotometer (Thermo Scientific, Madison, USA). Soil gravimetric water and organic matter were determined through soil drying at 105 °C for 12 hours for weight loss of ± 10 g followed by dry combustion of the organic material at 650 °C for 2 hours.

### Molecular analyses

DNA was extracted from 75 mg and 250 mg of dried roots and soil respectively using the Soil DNA Isolation Plus Kit (Norgen Biotek Corp) according to manufacturer’s instructions. Root samples were shredded using the Bead mill homogenizer (Bead Ruptor Elite, Omni International) prior to extraction. Amplification of the 18S SSU rDNA region of the extracted genomic DNA was done using AMF specific primer pairs (AMV4.5NF (forward primer) and AMDGR (reverse primer)) (Van Geel et al., 2014). The primers used were modified versions, including Illumina adaptors and indexes, for dual-index sequencing on Illumina MiSeq (Kozich et al. 2013). PCR amplification was carried out in 25 μL volumes comprising 1 µL of template DNA, 0.35 µL of each 10 µM primer, 12.5 µL ALLin HS Red Taq Mastermix (consisting of 0.25 mM dNTPS, 3 mM MgCl_2_, Hot Start Taq DNA Polymerase (2 u/µL), HighQu) and 10.8 µL PCR grade water. PCR reaction in Biometra TAdvanced Thermal Cycler 96 (Westburg Laboratories, Netherlands) started with 95 °C for 2 min (initial denaturation step), followed by 40 cycles of 15 s at 95 °C (denaturation), 15 s at 52 °C (annealing) and 15 s at 72 °C (elongation) and a final extension time of 5 min at 72 °C.

Purification of PCR products was done using Agencourt AMPure XP beads (Beckman Coulter, CA, USA), while DNA concentration and purity were measured using a Qubit fluorometer (Invitrogen). Samples were pooled in equimolar quantities. The quality of the amplicon libraries was assessed using an Agilent Bioanalyzer 2100 and high-sensitivity DNA chip (Agilent Technologies, Waldbronn, Germany). DNA fragments of the appropriate size (350 bp) were extracted from the gel and purified using QIAquick gel extraction kit (Qiagen). Lastly, sequencing of libraries was done at Genomics Core in Leuven, Belgium on the Illumina Miseq 2 × 250 platform.

### Bioinformatics

Demultiplexed reads of sequencing were merged, quality filtered and clustered into OTUs using USEARCH (Edgar 2010). First, forward and reverse reads were merged using the *fastq_mergepairs* command with a maximum of 10 mismatches and a minimum 80% id of alignment to the MaarjAM database. From these merged pairs, sequences shorter than 200 bp and with more than 1.0 expected errors were filtered out using the *fastq_filter* command. Unique sequences were identified and singletons were removed using the *fastx_uniques*. With the *‘cluster_otus’* command, sequences were clustered into operational taxonomic units (OTUs) at 97% sequence similarity threshold, and chimeric OTUs were removed. An Operational taxonomic Unit (OTU) table was then constructed using the *‘otutab’* command to assign a sequence to an OTU and a sample ID. In order to reduce the number of erroneous sequences produced during PCR and sequencing (Alberdi et al. 2018), an extra filtering step was performed by removing OTUs that were represented by less than 0.01 % of the sequences in a sample. For the taxonomic annotation, consensus sequence from OTUs were BLASTed against the MaarjAM 18s database (Öpik et al. 2010), and yet unidentified OTUs were further BLASTed against the National Centre for Biotechnology Information (NCBI) database with the following criteria for a hit: sequence similarity at least 97%, alignment at least 95% and E-value < 1^e-50^.

### Data analyses

Data obtained were analyzed in R version 4.3.2 (R Development Core Team 2021). Statistics on soil chemical properties were computed using Microsoft Excel 2010. R software packages such as vegan and ggplot2 were used for diversity and community composition analyses and graphical illustrations, respectively (Oksanen et al. 2022; Wickham 2016). Rarefaction curves were plotted to check whether the samples were sufficiently deep sequenced using the *rarecurve* command in R (Fig. S1a and b). OTU richness, Shannon diversity and Pielou’s evenness were calculated for each sample. Analysis of variance (ANOVA) was carried out to test if there was any difference in AM fungal diversity between the root and soil samples and between the studied tree species.

Further analyses were conducted for the ten tree species for which at least two sufficiently deep sequenced samples were available. These were *Albizia ferruginea* (*n* = 2), *Albizia zygia* (*n* = 2), *Amphimas pterocarpoides* (*n* = 3), *Caesalpinia bonduc* (*n* = 3), *Celtis zenkeri* (*n* = 3), *Entandrophragma candollei* (*n* = 4), *Lecaniodiscus cupanoides* (*n* = 4), *Mansonia altissima* (*n* = 3), *Pseudospondias microcarpa* (*n* = 2), *Triplochiton scleroxylon* (*n* = 2). This dataset contained 147 AM fungal OTUs. AM fungal community composition was visualized using principal components analyses (PCA) on the Hellinger-transformed OTU table. A permutational analysis of variance (PERMANOVA) was used to assess the effect of tree species on AM fungal community composition using the adonis function of the *vegan* package in R. This was repeated for a subset of the data excluding the tree species from the family Fabaceae, since these also associate with nitrogen-fixing bacteria and the tripartite interaction is known to affect AM fungal communities (Tsiknia et al. 2021). An indicator OTU analysis was conducted using the *indicspecies* package in R (De Caceres et al. 2016). Following this, we constructed a bipartite network to elucidate the architecture of the network of associations between trees and AM fungi. To get more insights into the structure of the plant-fungus network, we calculated the *H*_2_ metric of specialization, interaction evenness and nestedness using the ‘bipartite’ v.2.9 package (Dormann et al. 2008) in R. Nestedness was calculated using the nested overlap and decreasing fill (NODF) (Almeida-Neto et al. 2008) and weighted NODF measure proposed by Galeano et al. (2009). Both measures vary between 0 (random structure) and 100 (perfect nestedness). Additionally, we estimated connectance and network modularity using Newman and Girvan’s metric (Newman and Girvan 2002). The significance of the observed network measures was assessed by comparison with the 95 confidence intervals obtained from 1000 iterations of four different null models. To test the robustness of the results and to assess the effect of rare OTUs on network results, these analyses were repeated for a subset of the data including only the most abundant OTUs (i.e., OTUs with > 500 sequences present in at least 2 samples). All these analyses were performed using the package bipartite v. 2.9 (Dormann et al. 2008) in R.

## Results

### Tree species diversity and soil chemical properties

We identified 23 dominant tree species in the study area belonging to 11 different families: Malvaceae (5 spp.), Fabaceae (4 spp.) Anacardiaceae (2 spp.), Moraceae (2 spp.), Meliaceae (2 spp.), Sapindaceae (2 spp.), Sapotaceae (1 sp.), Myristicaceae (1 sp.), Annonaceae (1 sp.), Combretaceae (1 sp.) and Cannabaceae (1 sp.) (Tab. S1). Chemical analysis of the sampled rhizosphere soils showed that they had a moderately neutral pH (7.35 ± 0.42) and average available phosphorus (8.17 ± 3.41 mg/kg soil), ammonium (0.90 ± 0.49 mg/kg soil) and nitrate (4.23 ± 1.71 mg/kg soil) concentrations. Gravimetric water content and organic matter content were 22.49 % and 5.24 %, respectively (Tab. S2).

### Arbuscular mycorrhizal fungi community composition and distribution

Data from 63 samples comprising 40 root samples and 23 rhizosphere soil samples from 23 tree species were sequenced with AM fungal specific primers. Quality filtering and clustering of the 18S SSU database resulted in 6,251,966 sequences belonging to 710 OTUs (Supplementary Fig. S1a). Rarefaction curves were plotted to check whether the samples had been sufficiently deeply sequenced (Fig. S1). Based on these, we removed 10 samples with very low sequence numbers (< 1000 sequences). Eventually, 3,286,506 sequences belonging to 194 OTUs were identified as AM fungi. The majority of the AM fungal OTUs belonged to the family Glomeraceae (126 OTUs and 96.85 % of the sequences), while the remaining OTUs belonged to Gigasporaceae (38 OTUs and 1.53 % of the sequences), Acaulosporaceae (17 OTUs and 0.85 % of the sequences), Paraglomeraceae (3 OTUs and 0.05 % of the sequences), Diversisporaceae (2 OTUs and 0.22 % of the sequences) and Ambisporaceae (2 OTUs and 0.06 % of the sequences). Six AM fungal OTUs (0.43 % of the sequences) could not be identified to family level. Seven AM fungal genera were identified: *Glomus, Scutellospora, Acaulospora, Paraglomus, Funneliformis, Diversispora* and *Ambispora* (Fig. 2).

**Fig. 2.**
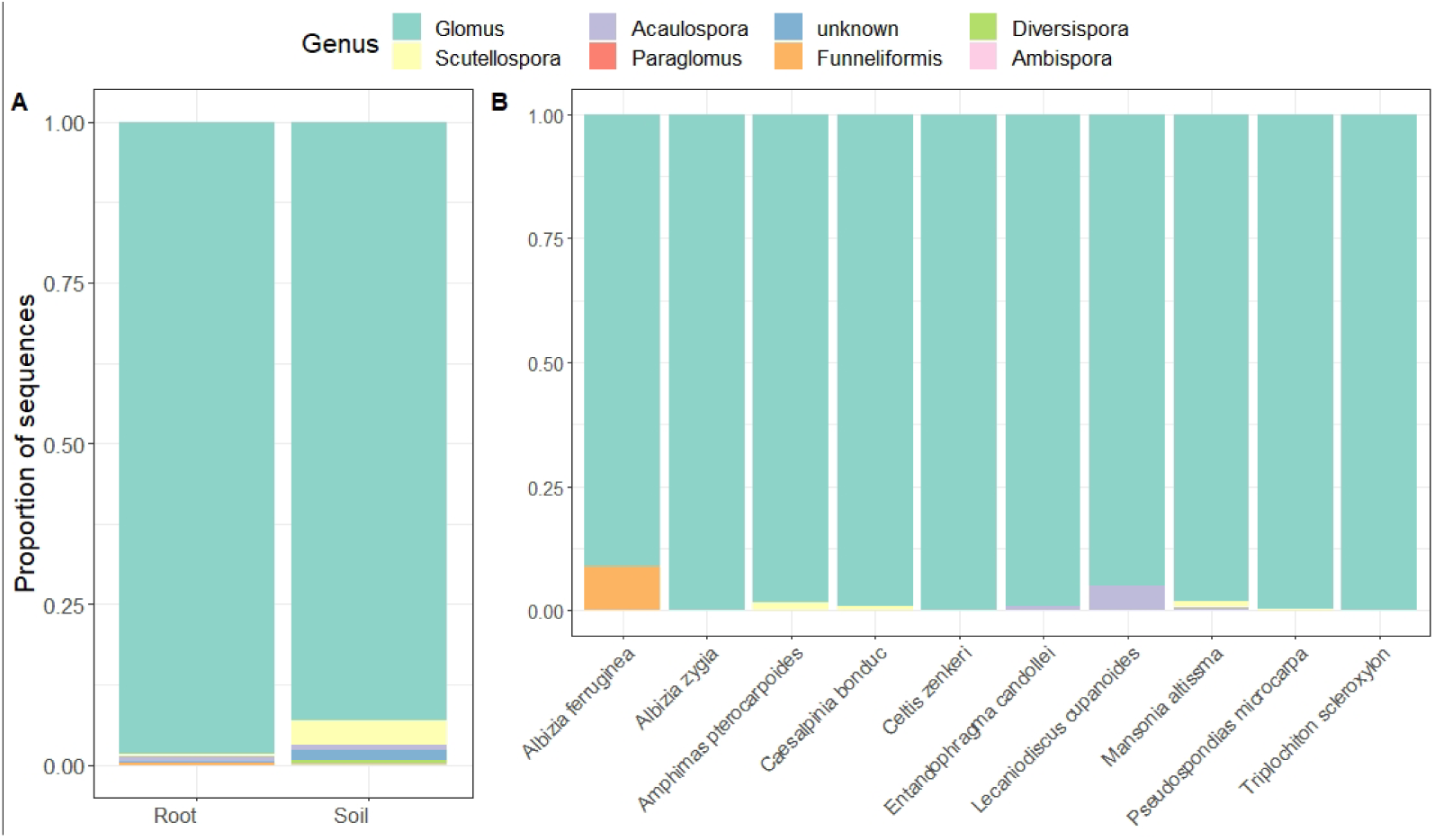
Frequency distribution of the sequences belonging to different AM fungal genera found associating with tropical rainforest trees in Nigeria, separate for (A) root (n = 30) and rhizospheric soil (n = 23) samples and (B) for the ten tree species for which there were at least two root samples left after quality filtering: *Albizia ferruginea* (n = 2), *Albizia zygia* (n = 2), *Amphimas pterocarpoides* (n = 3), *Caesalpinia bonduc* (n = 3), *Celtis zenkeri* (n = 3), *Entandrophragma candollei* (n = 4), *Lecaniodiscus cupanoides* (n = 4), *Mansonia altissma* (n = 3), *Pseudospondias microcarpa* (n = 2), *Triplochiton scleroxylon* (n = 2).

### AM fungal diversity analyses among tree species

We compared OTU richness, Shannon diversity and Pielou’s evenness between root samples and rhizosphere soil samples (Fig. 3A). The results showed that rhizosphere soil samples had significantly higher OTU richness (*F*_1,51_ = 32.34, *p* < 0.0001), Shannon diversity (*F*_1,51_ = 43.64, *p* < 0.0001) and Pielou’s evenness (*F*_1,51_ = 19.81, *p* < 0.0001). Root samples had on average (± standard deviation) 31 ± 12 OTUs, Shannon diversity 1.7 ± 0.5 and Pielou’s evenness 0.5 ± 0.1, while soil samples had on average 52 ± 15 OTUs, Shannon diversity 2.5 ± 0.4 and Pielou’s evenness 0.7 ± 0.1. None of the diversity indices differed significantly between the sampled tree species for which a sufficient number of samples was available (Fig. 3B). Ordination with principal components analysis (PCA) showed that, although there was some overlap, tree species did differ in their AM fungal communities (Fig. 4). Three of the four species belonging to the family Fabaceae (*Albizia zygia, Amphimas pterocarpoides* and *Caesalpinia bonduc*) clustered together. Permutation multivariate analysis of variance (PERMANOVA) confirmed that AM fungal communities significantly differed between the ten host tree species (*F_9,27_*= 2.42, *p* = 0.001, *R^2^* = 0.547). Excluding the four Fabaceae species did not change the result (*F_5,17_* = 2.45, *p* = 0.001, *R^2^* = 0.505). Five of the ten tree species had significant indicator OTUs associated with them (Table S3). *Entandrophragma candollei* had 6 indicator OTUs, five of which belonged to the genus *Acaulospora* and one to *Glomus. Albizia zygia* had two indicator OTUs, both belonging to the genus *Glomus. Amphimas pterocarpoides, Mansonia altissima* and *Triplochiton scleroxylon* each had one indicator OTU, belonging to respectively *Glomus, Scutellospora* and *Glomus*.

**Fig. 3.**
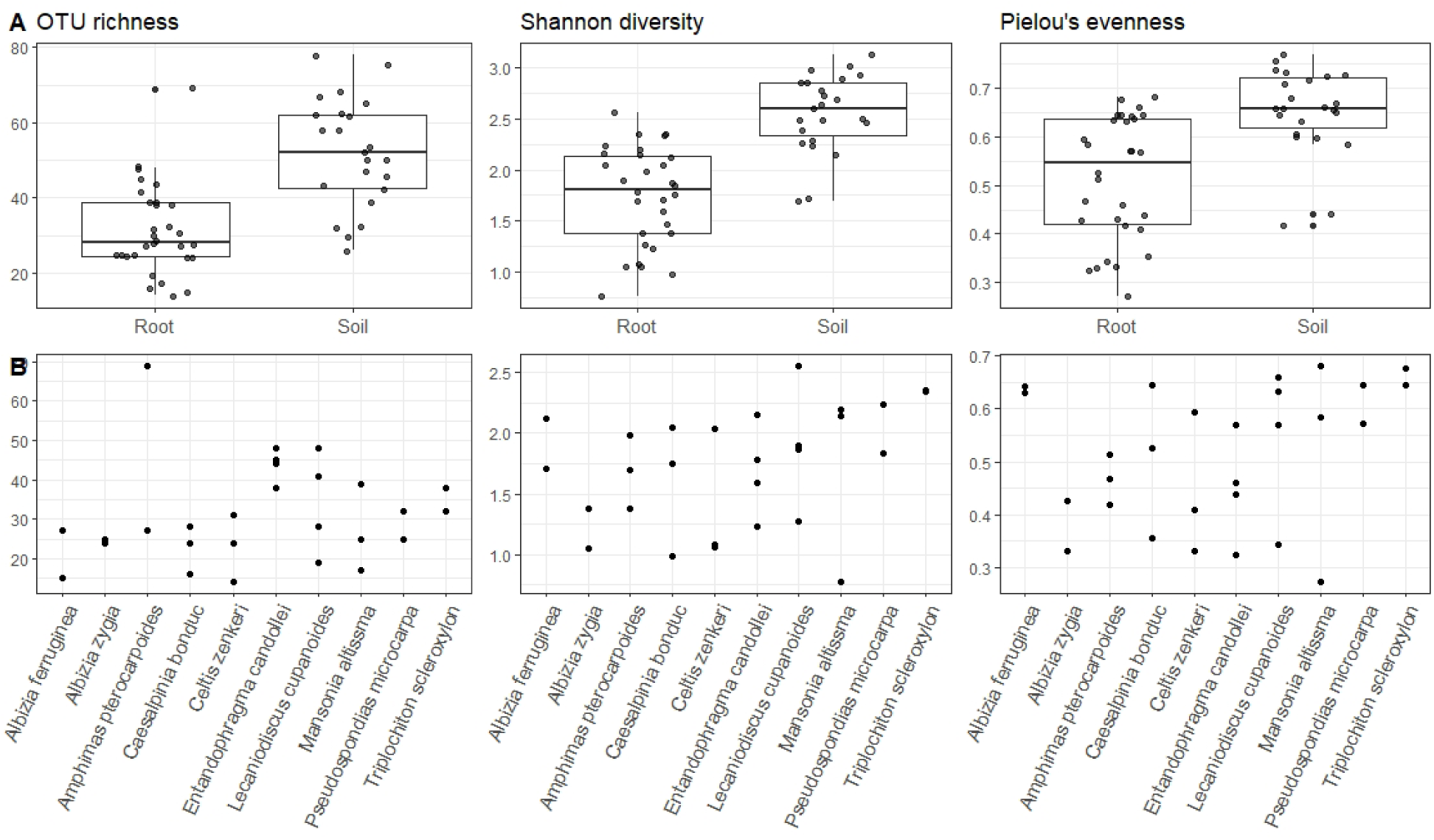
Differences in OTU richness, Shannon diversity and Pielou’s evenness between (A) root samples and rhizosphere soil samples and (B) tree species. None of the diversity indices differed significantly between the ten tree species for which at least two samples were sufficiently deep sequenced to be included in the analysis. For an overview of the diversity indices in all samples, see Fig. S2.

**Fig. 4.**
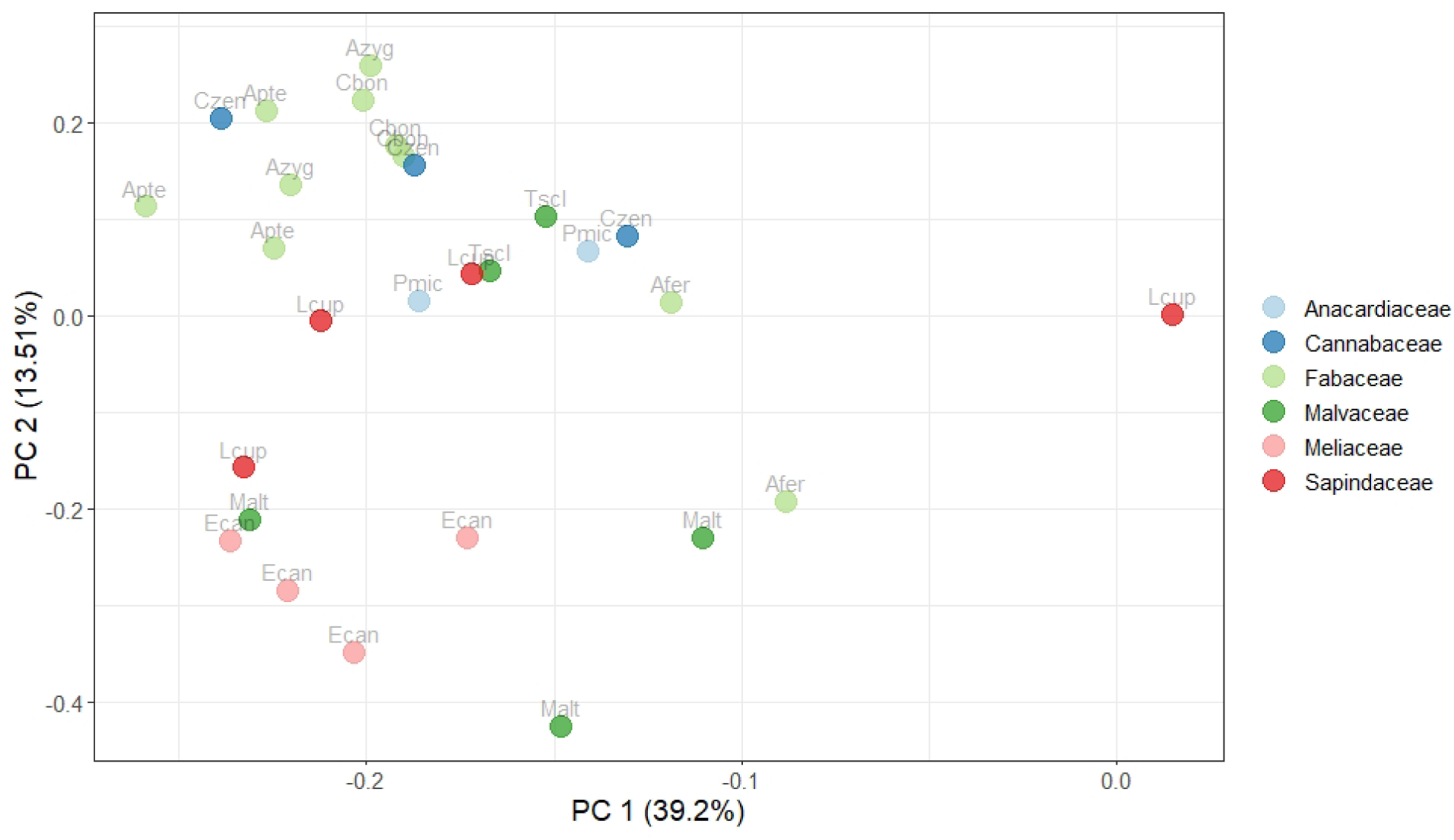
Principal components analysis showing the similarity between the root AM fungal communities of the ten species for which there were at least two root samples left after quality filtering: *Albizia ferruginea* (n = 2), *Albizia zygia* (n = 2), *Amphimas pterocarpoides* (n = 3), *Caesalpinia bonduc* (n = 3), *Celtis zenkeri* (n = 3), *Entandrophragma candollei* (n = 4), *Lecaniodiscus cupanoides* (n = 4), *Mansonia altissma* (n = 3), *Pseudospondias microcarpa* (n = 2), *Triplochiton scleroxylon* (n = 2). Each sample is colored according to the plant family to which the host tree belonged and marked with an abbreviation of the tree species name.

### Network analysis of AM fungal community among tree species

Network analyses showed that overall network connectance was 0.3626, meaning that 36.26 % of all possible interactions were observed. Trees shared on average 53.3 AM fungi, while AM fungi shared on average 3.6 trees (Fig. 5). The level of specialization *H*_2_ was 0.4272. The network was not significantly nested (NODF = 27.38, weighted NODF = 34.10, *p* > 0.05), but showed significant modularity (*M* = 0.4426). Six modules were detected, consisting of 3 species (*Albizia zygia, Amphimas pterocarpoides* and *Caesalpinia bonduc*), 2 species (*Celtis zenkeri* and *Lecaniodiscus cupanoides*), 2 species (*Entandrophragma candollei* and *Mansonia altissima*) and 3 times one species (*Albizia ferruginea, Triplochiton scleroxylon* and *Pseudospondias microcarpa*). Using a subset of the data, including only the most abundant OTUs, i.e. OTUs > 500 sequences, produced very similar results. In that case, *H*_2_ was 0.4260, NODF and weighted NODF were respectively 37.75 and 38.50 and *M* was 0.4435. Using this subset, four modules were detected, consisting of 2, 2, 4 and 2 species.

**Fig. 5.**
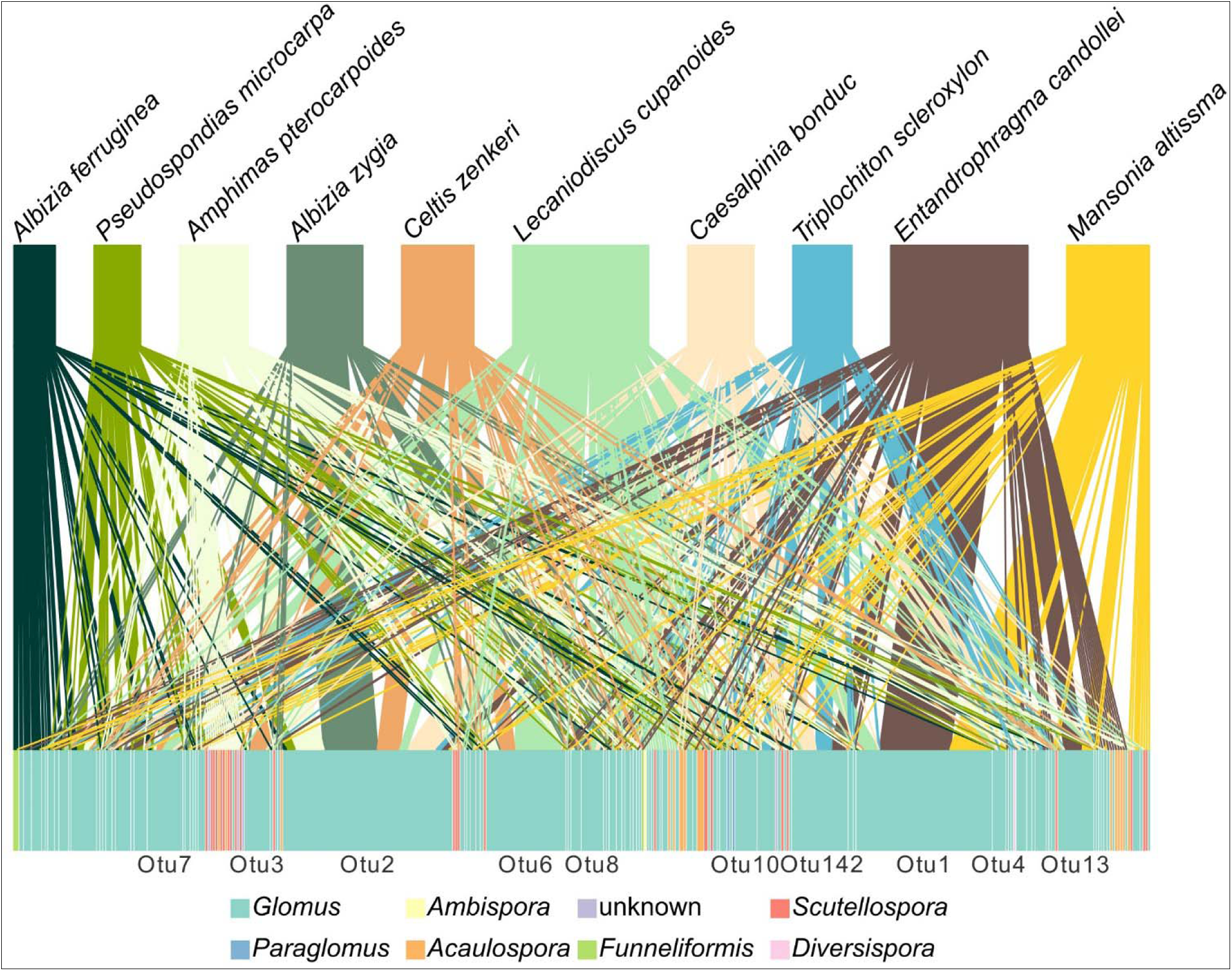
Visualization of the network of interactions between trees (upper boxes) and arbuscular mycorrhizal (AM) fungal operational taxonomic units (OTUs) (lower boxes).

## Discussion

In this study, we analyzed the AM fungi associated with tropical trees in African rainforest. We detected a total of 194 AM fungal OTUs associated with the studied host tree species. While some of the tree species have been microscopically examined before for the presence of mycorrhizal structures (Onguene and Kuyper 2001; Bechem et al. 2018), to our knowledge, this was the first time that their AM fungal communities were determined using molecular methods. The most abundant AM fungus genus was *Glomus*, aligning with previous studies on AM fungal composition in forest communities across the globe (Chen et al. 2017; Sheldrake et al. 2017, 2018; Boeraeve et al. 2019a; Ji et al. 2020; Touré et al. 2020; Suárez et al. 2023). The most abundant OTUs corresponded to taxa that have been found in (sub)tropical regions before. For example, VTX00069, the closest match in the MaarjAM database (Öpik et al. 2010) to our most abundant OTU, has been found in a tropical forest in Panama (Husband et al. 2002), savannah in Madagascar (Yamato et al. 2009), agricultural land in subtropical Mexico (Alguacil et al. 2008), a tropical mountain rainforest in Ecuador (Kottke et al. 2008), and a tropical forest in French Polynesia (Davison et al. 2018). The second most abundant OTU could not be matched with a virtual taxon in the MaarjAM database, but did match with a sequence in the GlobalAMFungi database (Větrovský et al. 2023), which came from a cocoa plantation in Côte d’Ivoire (Rincón et al. 2021). Also our third most abundant OTU matched with a virtual taxon from the MaarjAM database (VTX00132) and could be traced back to many of the aforementioned studies in (sub)tropical regions (Husband et al. 2002; Yamato et al. 2009; Davison et al. 2018), although it has also been observed in several studies in temperate or Mediterranean regions (de Leon et al. 2016; Davison et al. 2018). This suggests that AM fungi can be biased or even limited in their distribution towards a climatic zone, as has been shown before (Öpik et al. 2010; Pärtel et al. 2017). Given the strong sampling bias towards temperate areas (Větrovský et al. 2023), it is vital that sampling efforts in (sub)tropical areas are expanded in order to better understand AM fungal biodiversity patterns across spatial scales (Alexander and Selosse 2009).

### AM fungal communities in soil and root samples

Within a root system or even a single root, several AM fungi coexist and hence complex communities can be found. However, what explains final community structure remains unclear. Here, we compared AM fungal OTU richness between rhizosphere soil and root samples. Our results consistently showed that OTU richness, Shannon diversity and evenness were significantly higher in the rhizosphere soil samples than in the root samples, but did not differ significantly among tree species. Additionally, all tree species associated with a large number of AM fungi simultaneously, leading to complex multispecies assemblages within roots. These results are in accordance with previous research that has shown that OTU diversity of rhizosphere samples is generally higher than that found in roots (Sheldrake et al. 2018; Boeraeve et al. 2019b; Větrovský et al. 2023). This suggests that the regional pool of AM fungi from which the trees can select their partners, is larger than what eventually occurs in the roots and that the members of local communities are selected by passing through abiotic and biotic filters (Jumpponen and Egerton-Warburton 2005; Vályi et al. 2016). Abiotic filters refer to local habitat conditions that select only those AM fungal taxa that are adapted to the abiotic conditions in that habitat. Since most tropical forest soils are generally poor in nutrients, we believe that the observed high AM fungal richness and diversity were sustained by competition for the limited available nutrients. Indeed, several observational studies have shown that increased nutrient availability decreases AM fungal diversity (Van Geel et al. 2018; Ceulemans et al. 2019). Similarly, nutrition addition experiments have shown that addition of phosphorous and nitrogen reduced the AM fungal abundance in roots and soil, but responses were generally more pronounced for soil communities than root communities (Sheldrake et al. 2018).

Biotic filters refer to selective processes, in which a tree selects AM fungi that are most beneficial for growth and survival (Bever et al. 2009; Kiers et al. 2011). Because AM fungal community composition differed significantly between the studied tree species, host filtering may have contributed to the observed variation in mycorrhizal communities. Furthermore, we found that the AM fungal communities of three of the four Fabaceae tree species, known to simultaneously associate with nitrogen-fixing bacteria and AM fungi, to cluster together in the ordination and to form a module in the interaction network. This suggests that the tripartite interaction between trees, AM fungi and nitrogen-fixing bacteria induces some biotic filtering (Tsiknia et al. 2021). Trees associating with nitrogen-fixing bacteria have been found to alter AM fungal communities in the soil (Guisande-Collazo et al. 2016), although their impact can differ between species (Hoogmoed et al. 2014). It is thus likely that at least some nitrogen-fixing trees associate with a more specialized AM fungal community. As this potentially more specialized community could have affected the PERMANOVA results, we repeated the analysis without including the Fabaceae tree species and found that AM fungal communities still differed significantly between tree species.

Besides, successional patterns and temporal turnover in fungal associations (Ventre Lespiaucq et al. 2021) may explain why OTU diversity in roots is lower than that in associated rhizosphere. In this case, the AM fungal communities associated with seedlings differ from those associated with adult plants. Previous research in tropical trees has already shown strong shifts in the mycorrhizal communities over time, with AM fungal types that were dominant in newly germinated seedlings being almost entirely replaced by previously rare types in the surviving seedlings the following year (Husband et al. 2002). Since we only sampled roots from adult trees, it is likely that some of the OTUs found in the soil associate with developing seedlings, but do no longer occur in the roots of adult trees.

### Network analysis of the interactions between AM fungi and tropical rainforest trees

Our results further showed that most AM fungal species showed low to moderate host specificity (Sepp et al. 2019; Boeraeve et al. 2021) compared to their ectomycorrhizal counterparts, which tend to exhibit a preference for specific host plant species, genera or families (Lang et al. 2011; Molina and Horton 2015). As a result, the network structure of the studied tropical trees and their AM fungi was well interconnected and showed a low level of specialization, confirming earlier studies that have shown that AM fungi are not host-specific and show strong niche overlap (Toju et al. 2015; van der Heijden et al. 2015). In addition, the network was not significantly nested, but showed weak modularity. Several other studies have observed that networks between trees and AM fungi were not significantly nested. For example, Toju et al. (2015) showed that plant-fungus networks from a cool-temperate, warm-temperate and subtropical forest lacked a nested structure. However, unlike our observations, these networks showed a significant ‘antinested’ topology, meaning that the observed nestedness was significantly smaller than that expected under random plant-fungus associations. While these patterns are seldom seen in aboveground interaction networks, they appear to be more common in underground networks, although the precise mechanisms that explain this antinested pattern remain unclear (Toju et al. 2015).

## Conclusion

There have been very few studies that have used high-throughput sequencing to identify the fungi that associate with trees in African tropical forest ecosystems. In this study, we have successfully identified and characterized the AM fungi that associated with tropical trees in the Ise forest reserve in southwest Nigeria. The dominant AM fungal OTUs seemed to occur predominantly in (sub)tropical regions. The results showed that AM fungal communities differed between tree species in composition but not in diversity. The interaction network had a high connectance and a low level of specialization, with trees sharing on average 53.3 AM fungal OTUs, while AM fungi shared on average 3.6 tree species. Overall, these results indicate that tropical tree species, while differing to some extent in their AM fungal communities, associate with a wide variety of AM fungi and that fungi are frequently shared among trees.

## Acknowledgments

This research was supported by Coimbra Scholarship for Young African Researchers. We thank Kasper van Acker and Gerrit Peeters for help with the analyses.

