## Supplemental tables and figures for "Arbuscular mycorrhizal fungal diversity and association networks in African tropical rainforest trees"

**Supplementary Information**

**Table S1**: List of sampled tree species and families

| S/No | Tree species | Family |
| --- | --- | --- |
| 1. | *Albizia zygia* (DC) J. F. Macbr | Fabaceae |
| 2. | *Albizia ferruginea* (Guill & Perr.) | Fabaceae |
| 3. | *Amphimas pterocarpoides* (Harms) | Fabaceae |
| 4. | *Antiaris africana* Engl. | Moraceae |
| 5. | *Blighia sapida* K.D. Koenig | Sapindaceae |
| 6. | *Celtis zenkeri* Engl. | Cannabaceae |
| 7. | *Chrysophyllum delevoyii* De Wild. | Sapotaceae |
| 8. | *Cola milleni* K. Schum | Malvaceae |
| 9. | *Caesalpinia bonduc* (L) Roxb. | Fabaceae |
| 10. | *Entandrophragma candollei* (Harms) | Meliaceae |
| 11. | *Ficus mucuso* L. | Moraceae |
| 12. | *Lecaniodiscus cupanoides* Planch ex. Benth | Sapindaceae |
| 13. | *Mansonia altissma* (A. Chev) | Malvaceae |
| 14. | *Sterculia tragacantha* Lindl. | Malvaceae |
| 15. | *Sterculia rhinopetala* K. Schum | Malvaceae |
| 16. | *Spondias mombin* L. | Anacardiaceae |
| 17. | *Pycnanthus angolensis* (Welw.) Warb | Myristicaceae |
| 18. | *Khaya ivorensis* (A. Chev) | Meliaceae |
| 19. | *Pterygota macrocarpa* K. Schum | Malvaceae |
| 20. | *Triplochiton scleroxylon* K. Schum | Malvaceae |
| 21. | *Terminalia superba* L. | Combretaceae |
| 22. | *Pseudospondias macrocarpa* (A. Rich) Engl. | Anacardiaceae |
| 23. | *Xylopia aethiopica* (Dunal) A. Rich | Annonaceae |

**Table S2:** Soil chemical properties (n = 23)

| Soil variable | Mean ± Standard Deviation |
| --- | --- |
| pH | 7.35 ± 0.42 |
| Phosphorus (mg/kg soil) | 8.17 ± 3.41 |
| Ammonium (mg/kg soil) | 0.90 ± 0.49 |
| Nitrate (mg/kg soil) | 4.23 ± 1.71 |
| Gravimetric water content (%) | 22.49 ± 1.17 |
| Organic matter content (%) | 5.24 ± 0.26 |

**Table S3**: Results of the indicator species analysis.

|  | stat | p.value | Genus | Closest species/VT match (MaarjAM) |
| --- | --- | --- | --- | --- |
| *Albizia zygia* |  |  |  |  |
| Otu60 | 0.971 | 0.016 | *Glomus* | VTX00121 |
| Otu59 | 0.814 | 0.041 | *Glomus* | VTX00101 |
| *Amphimas pterocarpoides* | | | | |
| Otu580 | 0.816 | 0.042 | *Glomus* |  |
| *Entandrophragma candollei* | | | | |
| Otu543 | 1.000 | 0.002 | *Acaulospora* | VTX00024 |
| Otu182 | 1.000 | 0.002 | *Acaulospora* | VTX00024 |
| Otu597 | 1.000 | 0.002 | *Acaulospora* | VTX00024 |
| Otu233 | 0.866 | 0.019 | *Acaulospora* | *mellea* |
| Otu406 | 0.866 | 0.017 | *Acaulospora* | VTX00024 |
| Otu365 | 0.866 | 0.031 | *Glomus* | VTX00096 |
| *Mansonia altissma* | | | | |
| Otu223 | 0.816 | 0.043 | *Scutellospora* |  |
| *Triplochiton scleroxylon* | | | | |
| Otu288 | 0.891 | 0.009 | *Glomus* | VTX00248 |
| *Entandrophragma candollei + Lecaniodiscus cupanoides* | | | | |
| Otu461 | 0.833 | 0.045 | *Glomus* | VTX00387 |
| *Albizia ferruginea + Albizia zygia + Triplochiton scleroxylon* | | | | |
| Otu360 | 0.811 | 0.049 | *Glomus* | VTX00146 |
| *Albizia ferruginea + Lecaniodiscus cupanoides + Pseudospondias microcarpa* | | | | |
| Otu727 | 0.837 | 0.045 | *Glomus* |  |
| *Caesalpinia bonduc + Entandrophragma candollei + Triplochiton scleroxylon* | | | | |
| Otu30 | 0.930 | 0.003 | *Glomus* | VTX00166 |
| Otu762 | 0.878 | 0.007 | *Glomus* | VTX00397 |
| Otu91 | 0.816 | 0.042 | *Glomus* | VTX00159 |
| Otu500 | 0.816 | 0.039 | *Glomus* | VTX00166 |
| *Lecaniodiscus cupanoides + Pseudospondias microcarpa + Triplochiton scleroxylon* | | | | |
| Otu21 | 0.901 | 0.023 | *Glomus* | VTX00322 |
| *Albizia zygia + Entandrophragma candollei + Lecaniodiscus cupanoides + Pseudospondias microcarpa* | | | | |
| Otu154 | 0.85 | 0.044 | *Glomus* | VTX00322 |
| *Amphimas pterocarpoides + Caesalpinia bonduc + Entandrophragma candollei + Triplochiton scleroxylon* | | | | |
| Otu142 | 0.934 | 0.010 | *Glomus* | VTX00397 |
| Otu37 | 0.893 | 0.029 | *Glomus* | VTX00397 |
| *Albizia ferruginea + Albizia zygia + Amphimas pterocarpoides + Entandrophragma candollei + Pseudospondias microcarpa* | | | | |
| Otu18 | 0.877 | 0.009 | *Glomus* | VTX00268 |
| *Albizia zygia + Amphimas pterocarpoides + Entandrophragma candollei + Mansonia altissma + Triplochiton scleroxylon* | | | | |
| Otu31 | 0.844 | 0.046 | *Glomus* | VTX00092 |
| *Albizia ferruginea + Amphimas pterocarpoides + Caesalpinia bonduc + Celtis zenkeri + Lecaniodiscus cupanoides + Mansonia altissma + Triplochiton scleroxylon* | | | | |
| Otu10 | 0.949 | 0.01 | *Glomus* |  |

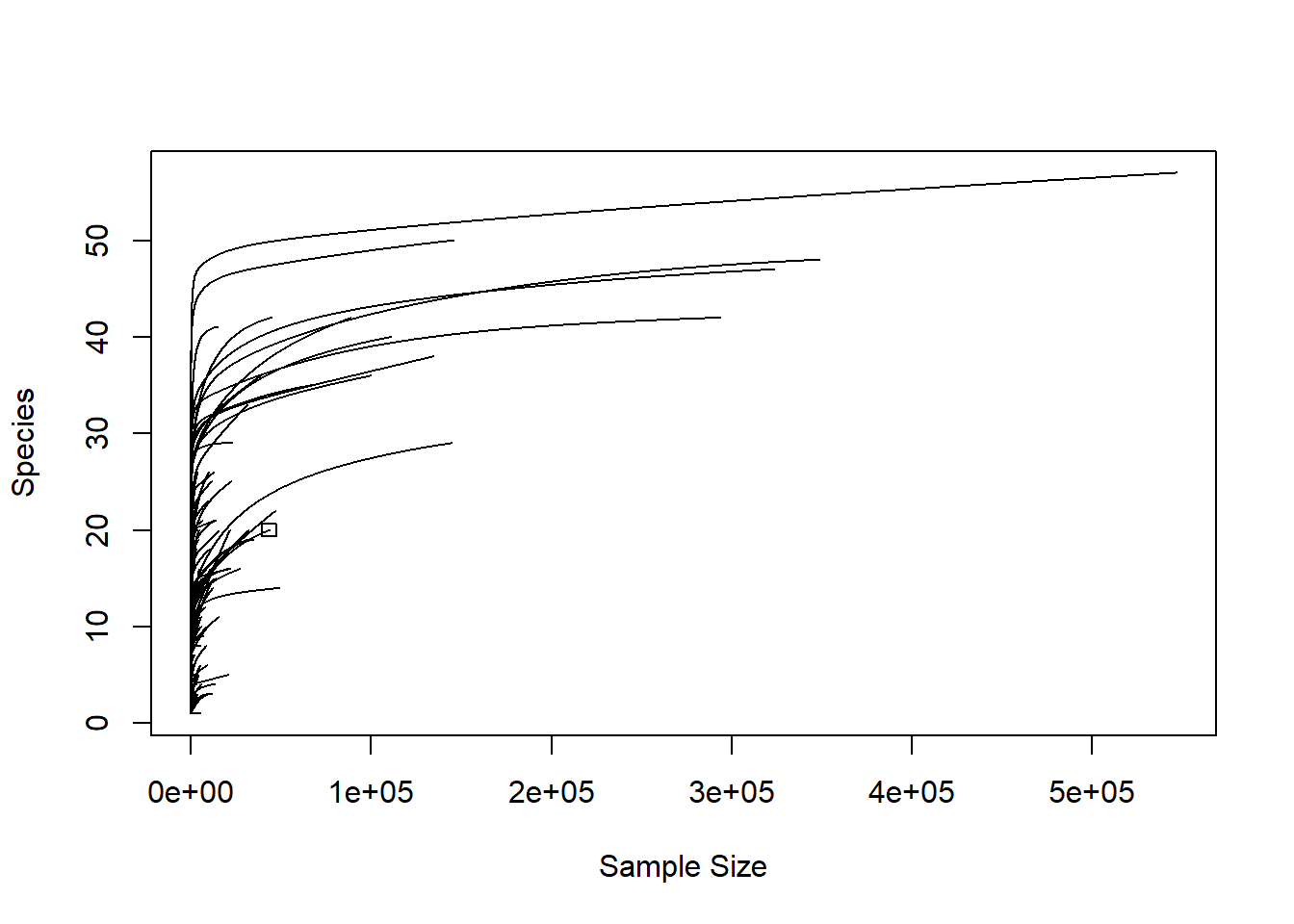

**Fig. S1a**: Rarefaction curve of all root and rhizosphere samples with insufficiently deep sequenced OTUs

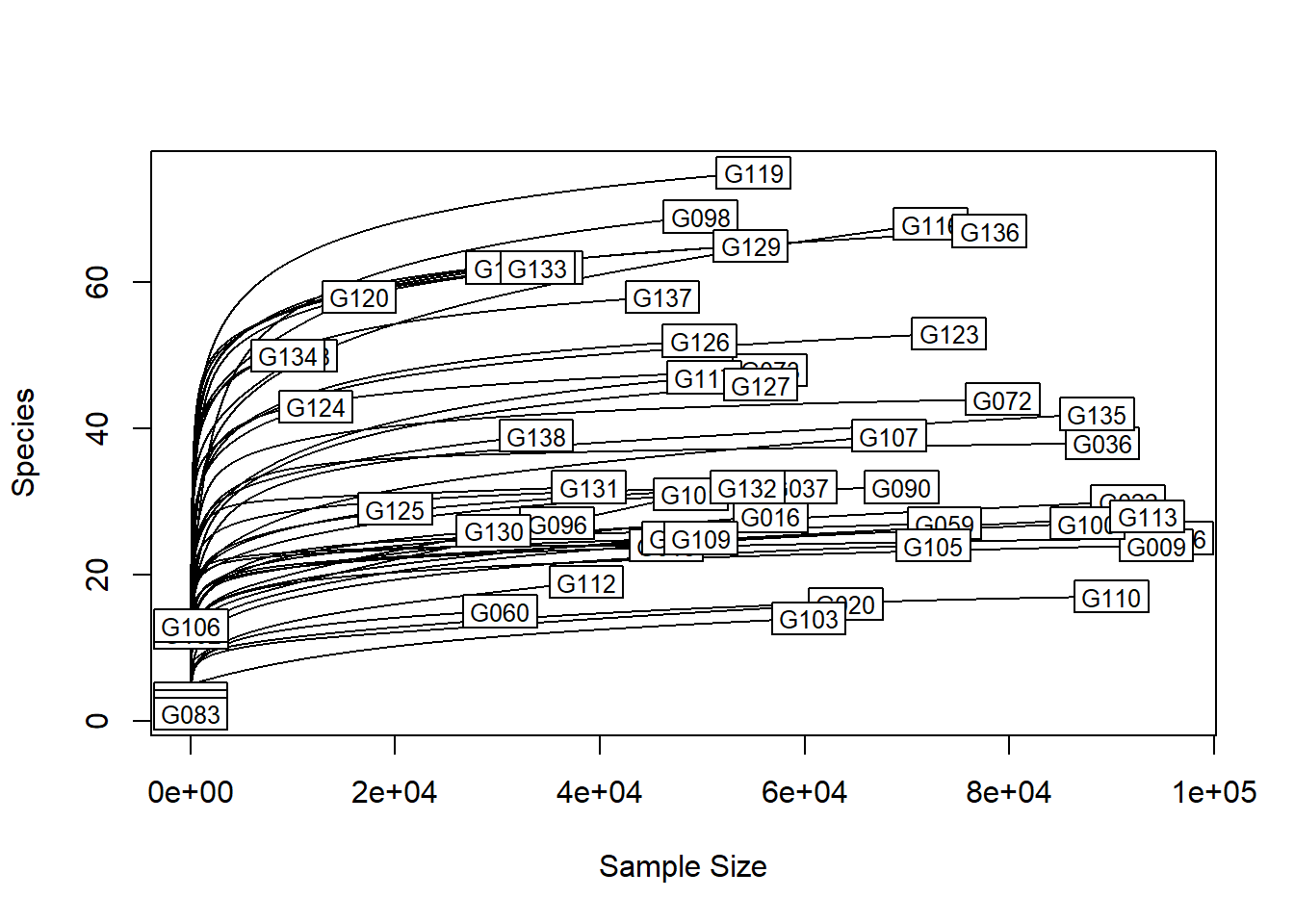

**Fig. S1b**: Rarefaction curve of retained samples upon removal of the insufficiently deep sequenced samples.

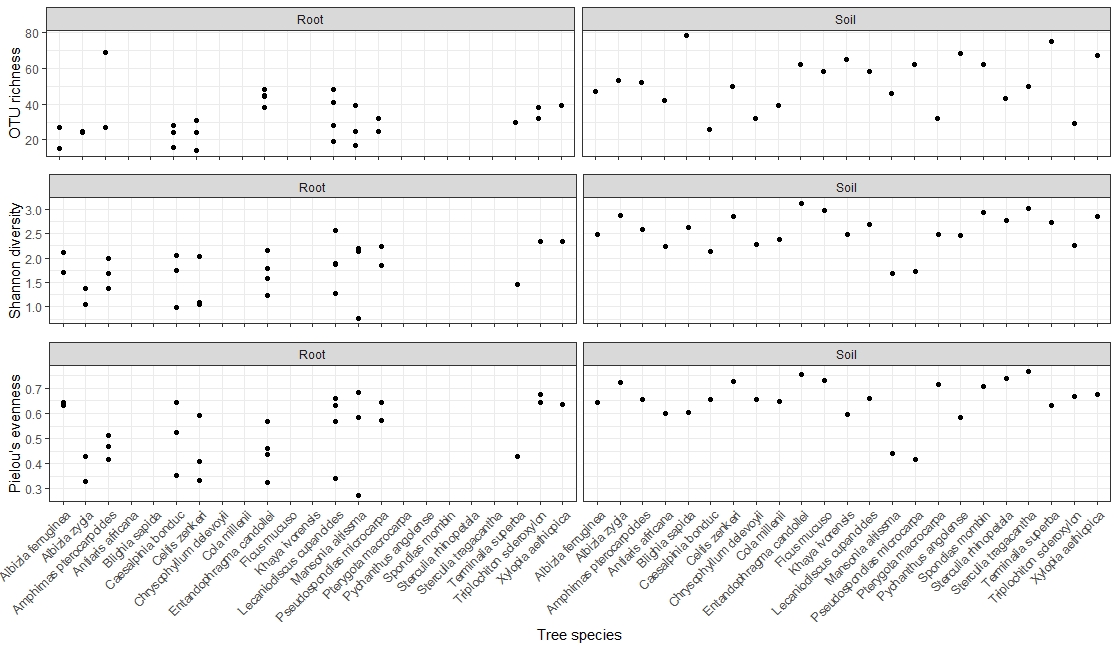
**Fig. S2** Diversity metrics per tree species, for the root and for the rhizospheric soil samples. Only samples that were sufficiently deep sequenced (i.e. > 10)
